# Symbiont virulence is a poor predictor of impacts on host population dynamics

**DOI:** 10.1101/2025.04.10.648206

**Authors:** Marcin K. Dziuba, Kristina M. McIntire, Elizabeth S. Davenport, Fiona E. Corcoran, Taleah Nelson, Paige McCreadie, Riley T. Manuel, Emma Baird, Natalia Ferreira dos Santos, Mia Robbins, Emma Dismondy, Kira J. Monell, Cristian Huerta, Lindsey C. Selter, Katya Deckelbaum, Michael H. Cortez, Meghan A. Duffy

**Author notes:** Hawai‘i Cooperative Studies Unit, University of Hawai‘i; Hilo, Hawaii, 96720, USA. Department of Ecology & Evolutionary Biology, Princeton University; Princeton, New Jersey, 08544, USA. Department of Epigenetics, Van Andel Institute; Grand Rapids, Michigan, 49503, USA. Animal Health Diagnostic Center, College of Veterinary Medicine, Cornell University; Ithaca, New York, 14850, USA.

## Abstract

Symbionts are classified as parasites or mutualists based on their impacts on host individuals, but timescale, ecological context, and level of biological organization can all influence symbiosis outcomes, and hence whether they are deemed harmful or beneficial. We designed experiments exposing the key freshwater grazer *Daphnia dentifera* to two symbionts varying in virulence, to test how well individual-level effects translate into population-level patterns. We found that a seemingly mildly virulent microsporidian strongly reduced host population size, with consequences for primary production, while a yeast that was highly virulent to individuals had no detectable negative effects on host population size. Our experiments show that studies that fail to consider the impacts of symbionts at multiple scales risk focusing mitigation and control measures on the wrong symbionts.

## Main text

Scientists have long recognized that symbionts can have major effects on host fitness (*1*), classifying symbionts with net positive impacts as mutualists and those with net negative impacts as parasites (*2*). This classification is typically based on individual-level measurements of changes in fitness-related traits of the host, like symbiont-induced changes in host survival (*3–5*) or reproductive success (e.g., (*5–7*)). A common measure of the impact of symbionts on hosts is virulence, a term that has a variety of definitions, but in a broad sense is defined as damage inflicted on the host by the symbiont (e.g., (*5*)), and in a narrow sense as symbiont-induced increases in mortality (*3–5, 8*). Here, we use the broad sense definition, but emphasize that both definitions of virulence refer to an individual-level measure of symbiosis outcome. Measuring virulent effects of parasites on life-history traits helps assess the impact of a given symbiont on its host, and is commonly used in human and veterinary medicine, conservation, and organismal biology (*4, 9, 10*). However, besides the individual-level impacts, it is also important to identify how symbionts affect host population trajectories; after all, a common reason for measuring individual-level effects is to predict the population-level patterns. Accurate prediction of the large-scale, long-term consequences of symbiosis outcomes is crucial to identify risks and benefits of symbiont spread through host populations, and can be used in improving management practices in agriculture and conservation biology (*11, 12*). Yet, there is growing evidence that effects at the individual-level can poorly predict effects at the population-level (*12*), and that symbionts that seem benign at the individual-level can have harmful population-level effects and vice versa (e.g., (*13, 14*)). These contradictory patterns stem from scale-dependent impacts like density-dependent effects, trans- and multigenerational effects or environmental context (*13, 15–18*), and studies of symbionts at the individual level cannot detect these impacts. Hence the individual-level studies of virulence are burdened with a risk of incorrectly classifying a symbiont as harmful, neutral or beneficial for the population. Such incorrect labeling of a symbiont can lead us astray when we attempt to assess the food web- and ecosystem-level impacts of symbionts, or when we try to decide which symbionts should be combated or propagated. To improve our understanding of the role symbionts play in host populations, food webs and ecosystems, we need to recognize that the outcomes of symbiosis can vary depending on the time scale, level of biological organization, and ecological context in which the outcome is measured (e.g., (*12–14*)).

In this study, we explored how the net impact of symbiosis may change with scale and context, using the freshwater zooplankton *Daphnia dentifera* as a model organism. *Daphnia*, like all organisms, are subject to and dependent on symbiotic interactions; some of these interactions are vital for their survival (e.g., *Daphnia* without gut microbiomes die (*19*)), while others are detrimental (e.g., highly virulent parasites, (*20*)). A well-studied symbiont of *Daphnia* spp. is *Metschnikowia bicuspidata*, a yeast pathogen considered highly virulent mostly due to its large reduction in host lifespan (*21–23*). A recently discovered and seemingly much less virulent microsporidian symbiont of *Daphnia* species, *Ordospora pajunii* (*24, 25*), shows potential to become a mutualist when *M. bicuspidata* is present in the environment (*26*). *Ordospora pajunii* infection reduces the penetrability of *Daphnia* guts to attacking *M. bicuspidata* spores (*26, 27*). Additionally, an experimental investigation of potential drivers of context-dependent mutualism revealed that *O. pajunii* causes *Daphnia* to become a dead-end host for *M. bicuspidata* upon sequential co-infection due to increased host mortality (*27*). However, *O. pajunii* has also been found to greatly reduce offspring quality through transgenerational effects, which should decrease host population size over longer time scales (*28*). Therefore, our expectation was that the net impact of *O. pajunii* might change depending on whether the impacts were measured at the individual-level over short timescales versus at the population-level over longer time scales, as well as depending on whether the ecological context included the highly virulent *M. bicuspidata*. We first investigated the short-term, individual-scale effects of each symbiont on the host with a life-table experiment, then tested for potential shifts in outcomes based on ecological context and longer timescales (spanning generations) using a population-level experiment.

### Individual-level effects

At the individual-level, infections affected host reproduction (parasite:clutch interaction *F*=43.52, *p*<0.001) and lifespan (parasite effect *F*=29.35, *p*<0.001), but the effects of each symbiont strongly differed. Specifically, the yeast *M. bicuspidata* was highly virulent to the host, reducing its reproduction by 28-46% in clutches 2-4 (*t*-ratio>5.4 and *p*<0.001 for each comparison, Fig. 1A) and lifespan by 63.5% (*t*-ratio=6.22, *p*<0.001, Fig. 1B), the latter resulting in the host producing no more than four clutches. *Ordospora pajunii* infection, on the other hand, was less harmful to host individuals. The infections of *O. pajunii* had no effect on the number of offspring in the first three clutches, but this microsporidian had a negative effect on later reproduction, with the size of clutches 4-8 reduced by 8-29% (*t*-ratio>3.2 and *p*<0.003 for each comparison, Fig. 1A) and clutches 9-14 reduced by 31-54% (*t*-ratio>11.2 and *p*<0.001 for each comparison). *O. pajunii* did not significantly reduce host lifespan (*t*-ratio=1.74, *p*<0.27, Fig. 1B). Parasite treatment had a significant effect on reproduction time of *Daphnia* in clutches 1-4 (parasite effect *F*=5.00, *p*=0.007) (Fig. S1). While neither *M. bicuspidata* nor *O. pajunii* exposed *Daphnia* differed from the control (C-MB *t*-ratio=-0.74 and *p*=1.000; C-OP *t*-ratio=1.43 and *p*=0.466), the *O. pajunii* exposed *Daphnia* reproduced faster than those exposed to *M. bicuspidata* (MB-OP *t*-ratio=3.15 and *p*=0.005) (Fig. S1). Overall, *M. bicuspidata* had larger negative effects on the reproduction and survival of individuals than *O. pajunii*.

**Fig. 1.**
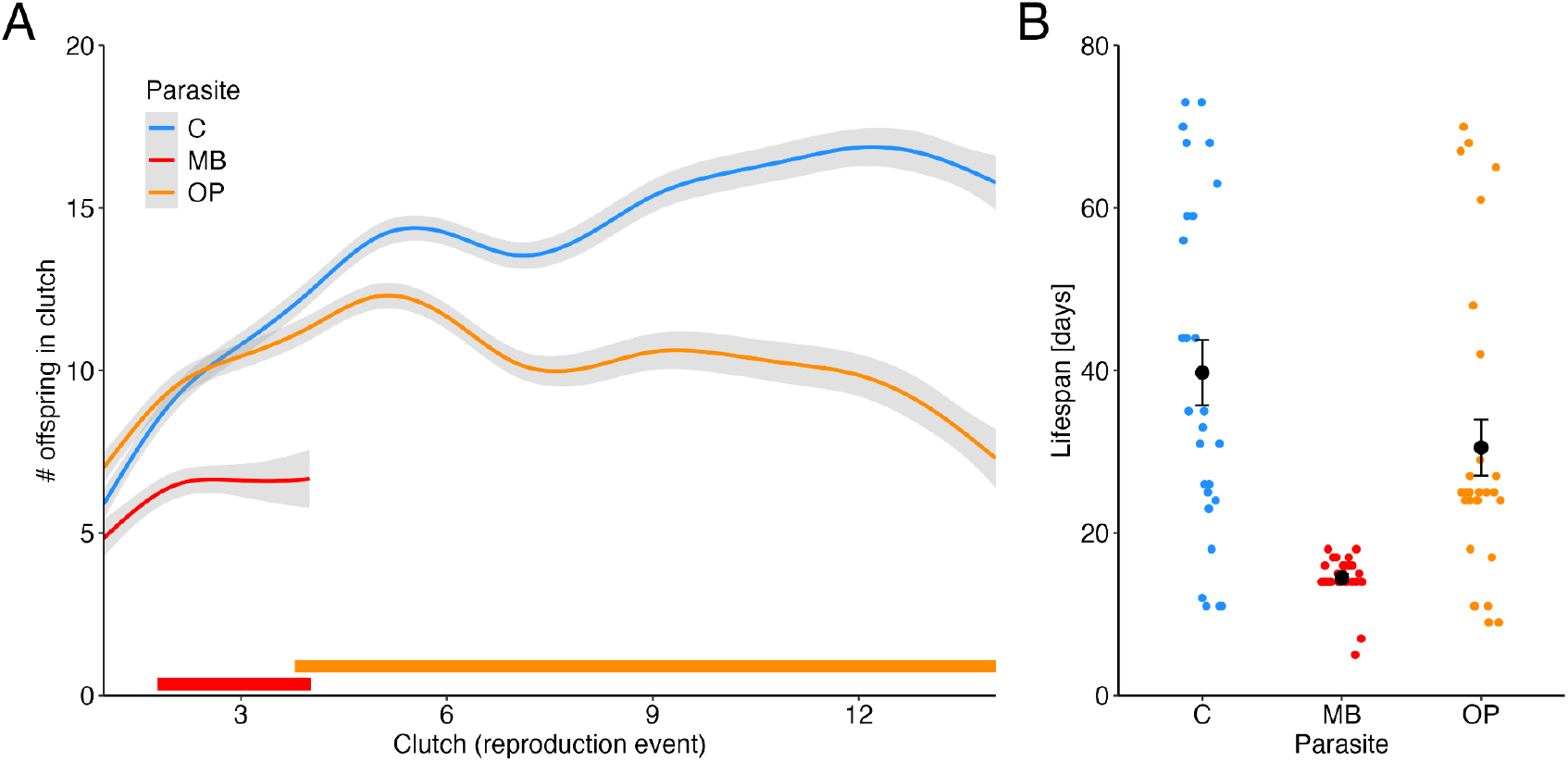
The number of offspring per clutch (A) and the lifespan (B) of *Daphnia* were strongly reduced by *M. bicuspidata* (red), while *O. pajunii* (orange) reduced clutch size only in late life stages (after clutch 3) and had no effect on host lifespan. Panel A shows fitted values±SE of clutch size (number of offspring per reproduction event) in consecutive clutches. Thick lines on the bottom of the figure indicate clutches at which the difference in clutch size between control and the treatment of corresponding color is significant. Panel B shows mean±SE lifespan (black points and error bars) of *Daphnia* in each treatment, with individual data points color-coded by treatment.

### Scaling up

Based on the life-table experiment results and previous studies of these symbionts, we predicted that the highly virulent yeast *M. bicuspidata* would strongly reduce the population density of *Daphnia*. The life-table experiment suggested that *O. pajunii* should have negligible effects on *Daphnia* populations. However, we know that *O. pajunii* induces negative transgenerational effects (i.e., transgenerational virulence, (*28*)), and therefore, we expected that the microsporidian could negatively impact *Daphnia* population density over multiple generations. We also hypothesized that *O. pajunii* would prevent, or at least reduce, *M. bicuspidata* outbreaks, benefiting the host. Hence, we exposed *Daphnia* populations to *O. pajunii, M. bicuspidata*, both, or neither as a control. Because *O. pajunii* outbreaks in natural lakes occur before those of *M. bicuspidata* (*27*), and the hypothesized protection mechanism against *M. bicuspidata* requires established infection of *O. pajunii* (*26, 27*), we added *O. pajunii* spores at the start of the experiment and waited until day 31 for its epidemics to be established at ∼15% prevalence to add *M. bicuspidata*. We analyzed the growth rates and densities of host populations, as well as prevalences of each parasite. We also investigated differences in host demographics, expecting that transgenerational virulence of *O. pajunii* would reduce the proportion of juveniles in the populations. Additionally, we quantified the standing stock of algae in the microcosms to determine how the symbionts altered host-resource interactions.

For the population-level experiment, the effects of single- and co-exposure to the symbionts were assessed during two time intervals: the initial growth phase (until day 23) and the post-peak phase (days 30-93). The separation was determined by both population dynamics and the experimental design: *M. bicuspidata* spores were added on day 31, when *O. pajunii* epidemics were established (∼15% prevalence) and the host populations had peaked in abundance. Thus, we used the time before the *M. bicuspidata* exposure to quantify how *O. pajunii* affects initial host population growth rate, and we combined the two groups exposed to *O. pajunii* into one group (OP and OPMB, labeled ‘OP’ in Fig. 2A), and the two unexposed groups into a separate group (C and MB, labeled ‘C’ in Fig. 2A). The post-peak phase was used to analyze the impact of both symbionts on host population trajectories, with the four treatments analyzed separately.

**Fig. 2.**
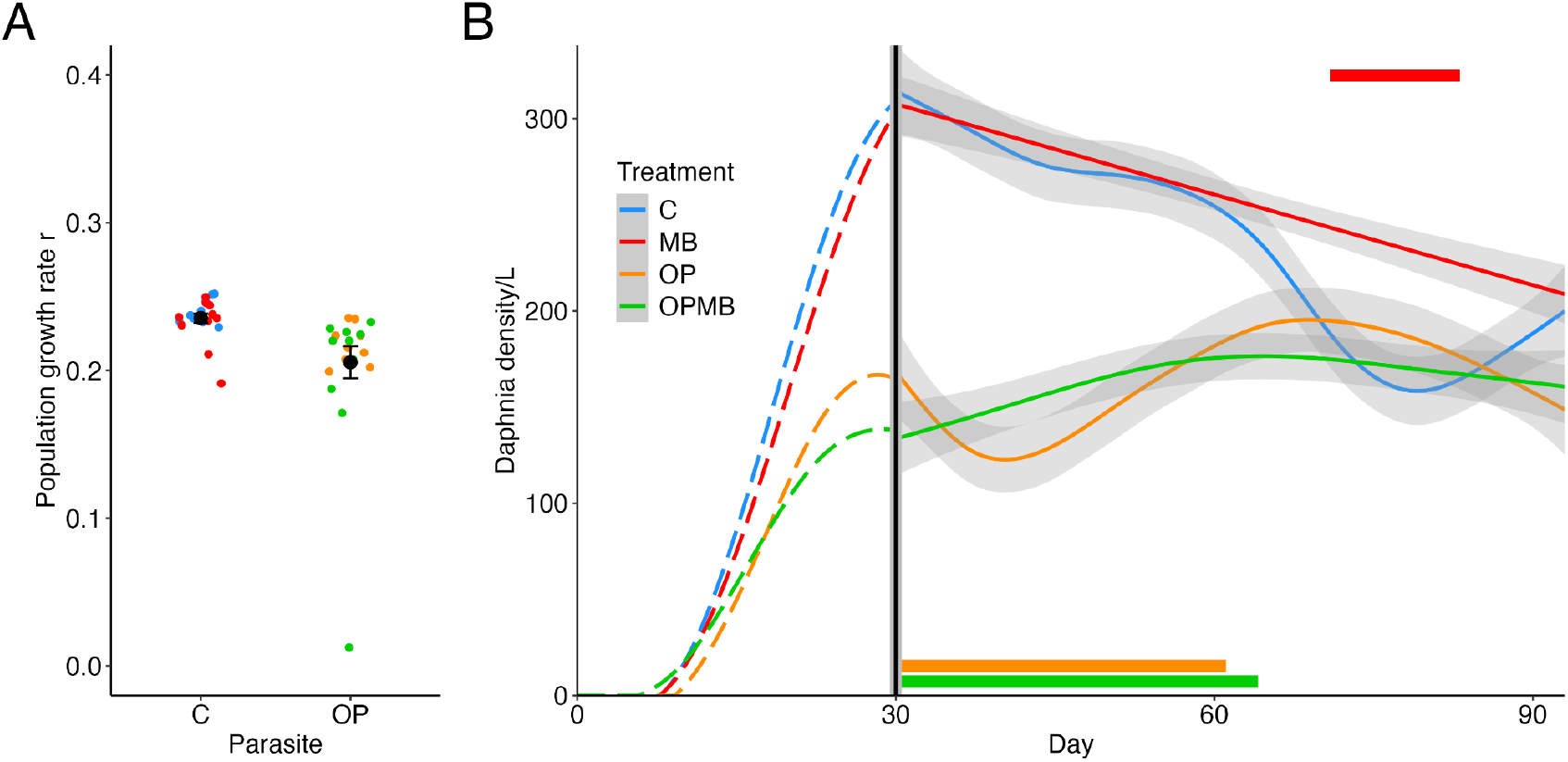
*Daphnia* initial population growth rate (A) and population size (B) were reduced by *O. pajunii* whereas *M. bicuspidata* increased population size. Panel A shows population growth rate during the initial growth phase (through day 23) in groups unexposed (C and MB) and exposed to *O. pajunii* (OP and OPBM), grouped into parasite treatment C or OP, respectively, because the population growth rates were measured before *M. bicuspidata* dosing. The black points and error bars indicate mean±SE, the data points are plotted underneath with colors corresponding to treatments. Panel B shows GAM modeled population densities in each treatment, the vertical gray line with black stripe on day 30 separates the time before the populations peaked (dashed lines) and after they peaked (solid lines); *M. bicuspidata* was dosed to the system on day 31, around when densities peaked. Only the modeled population sizes after day 30 were analyzed with the GAM. The population size in the single infection (OP) treatment was reduced on days 30-61, while, in the co-infection (OPMB) treatment, it was reduced on days 30-64. *M. bicuspidata* (MB) had a positive effect on the host population on days 71-83. The solid lines and gray ribbons indicate fitted values±SE, thick lines on top and bottom of the figure indicate the time span at which the treatment of the corresponding color had significantly larger or smaller host population size, respectively, in comparison to the control.

### Population-level effects

Contrary to what the individual-level measurements of virulence predicted, we found strong negative effects of *O. pajunii* on the host populations and no negative effects of *M. bicuspidata*. In the initial growth phase of the experiment, *O. pajunii* reduced the population growth rate by 13% (*χ*^2^ = 17.37, *p*<0.001, Fig. 2A). This negative impact of *O. pajunii* on hosts continued after the peak in densities. In populations with only *O. pajunii*, host population size on days 30-61 was reduced by, on average, 47% (*t*-ratio>2.7 and *p*<0.034 for each comparison between OP and C on each day); in the co-exposure treatment, host population size on days 30-64 was reduced by, on average, 42% (*t*-ratio>2.8 and *p*<0.026 for each comparison; Fig. 2B). Conversely, *M. bicuspidata* had no observable negative effects on the host populations; instead, later in the experiment (days 71-83), the *Daphnia* population size was larger in comparison to the control populations by, on average, 41% (*t*-ratio<-2.84 and *p*<0.028 for comparisons on each day; Fig. 2B).

The most likely explanation for the decimation of *Daphnia* populations by *O. pajunii* is transgenerational virulence (i.e., negative effects of maternal exposure on offspring fitness, even when those offspring are not themselves exposed (*28*)). A mathematical model estimating the population-level impacts of transgenerational virulence predicted an approximately 30% reduction in population size (*28*), which is similar to the effect we observed in the population experiment. Additionally, we found a lower proportion of juveniles in the *O. pajunii-*exposed populations (treatment effect *F*=9.66, *p*<0.001; Fig. 3), which is consistent with high juvenile mortality caused by transgenerational virulence. Adverse transgenerational effects of maternal exposure to stressors have been observed before, via, for example, altered embryo nutrition or epigenetic inheritance (*29, 30*) or, for some parasites, vertical transmission (*31*). We do not know yet the process responsible for the transgenerational virulence of *O. pajunii*, but we do know that the microsporidian is not vertically transmitted and that the embryos produced by exposed mothers are more fragile (*28*).

**Fig. 3.**
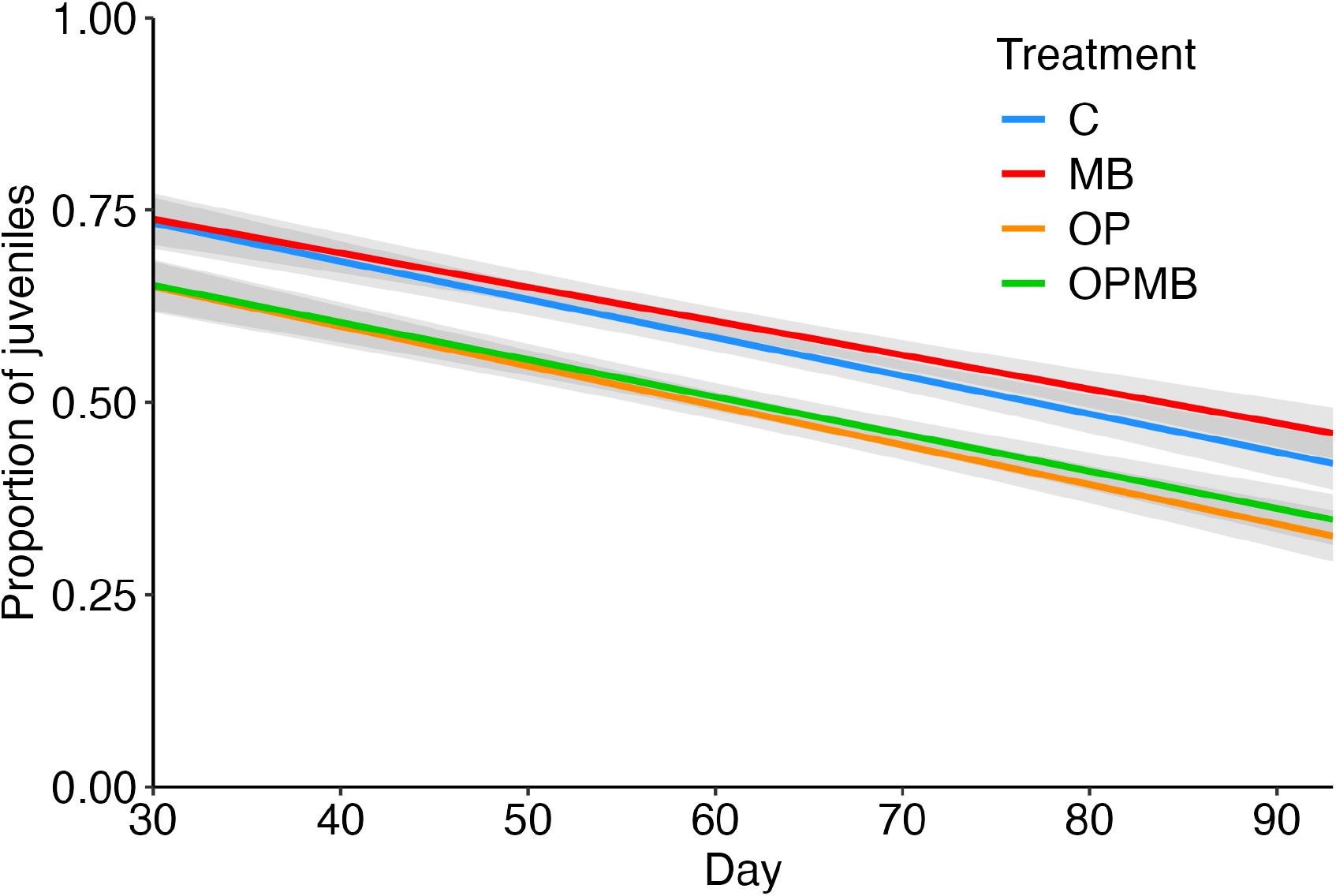
Proportion of juveniles decreased over time in all treatments, but populations with *O. pajunii* outbreaks consistently had a lower proportion of juveniles than either control populations or *M. bicuspidata* exposed populations. Fitted values of linear regression±SE are presented.

### Resource availability

We also hypothesized that some of the discrepancy between the individual-level effects on host fitness and population-level patterns could be explained by host-resource interactions. Some models indicate that intraspecific host competition for resources can amplify the virulence of seemingly benign parasites (*13*). Specifically, strong negative effects of *O. pajunii* could be the result of an infection-driven increase in resource demand that led to starvation and smaller population sizes. Contrary to this, *O. pajunii* exposed populations had leftover algae on days 29-56 in single exposure and on days 25-64 in co-exposure (C-OP *t*-ratio<-2.88 and *p*<0.025 ; C-OPMB *t*-ratio<-2.68 and *p*<0.045 for the given time spans) (Fig. S2), when the host populations were undergoing the strongest reductions in density. This indicates that the decrease in population size of *Daphnia* exposed to the microsporidian was not driven by resource depletion and starvation. Conversely, it is possible that the exposed *Daphnia* were undergoing illness-mediated anorexia (*32, 33*), which would lower the overall food consumption. In a different study, we found that *O. pajunii* can depress host feeding rates (*34*). Hence, the unexpected harmfulness of *O. pajunii* is not likely to be a result of resource shortage. We acknowledge that *O. pajunii* infection might hamper host nutrition through processes independent of food density in the environment, although assimilation efficiency of hosts seems to be unaffected by the microsporidian (*34*).

In contrast, *M. bicuspidata-*exposed *Daphnia* populations consumed all the available food (Fig. S2). Previous work has suggested that infection-driven decreases in foraging rates in *M. bicuspidata*-infected hosts (*32, 35*) can allow for increased resource uptake by other (susceptible, uninfected) hosts, driving a phenomenon known as a disease-induced hydra effect (*36*). The lack of a standing stock of algae in the *M. bicuspidata-*exposed treatment suggests that the very high densities of hosts consumed all the available food; if infected hosts consumed less food, that would leave additional food for the uninfected hosts, fueling a hydra effect, which is consistent with the periodically higher population size of the yeast-exposed population in comparison to the control (Fig. 2B). Additionally, during *M. bicuspidata* epidemics, the older and larger *Daphnia* experience the greatest infection-driven mortality. Old and large adults are effective grazers (*37*), but contribute very little to the population growth rate (*38*). Therefore, their increased mortality would release a fraction of resources from their control, and nourish the juveniles, increasing their survival, and stabilizing the whole population. Hence, we suspect that a joint effect of increased resource availability due to foraging depression, and age-dependent mortality eliminating large grazers, could be responsible for the lack of negative effects of *M. bicuspidata* at the population-level despite its high individual-level virulence.

### Antagonistic relationship of the symbionts

This study was motivated by the hypothesis that *O. pajunii* is a context-dependent mutualist whose impact switches from mildly negative to positive in the presence of the more harmful yeast symbiont. Consistent with prior work on *O. pajunii* (*26, 27*) and on its close relative (*39*), *M. bicuspidata* prevalence was reduced in our experiment in the presence of *O. pajunii*, and *O. pajunii* prevalence was lower during *M. bicuspidata* outbreaks (Fig. 4A). Specifically, the prevalence of *M. bicuspidata* was lower in co-exposure from day 56 of the experiment onward, and reached a 66% reduction by the end of the experiment (treatment:day interaction *z*-value=-2.33, *p*=0.020; Fig. 4A). *Ordospora pajunii* was also less prevalent in the co-exposure treatment from day 44 onward, reaching a 57% reduction at the end of the experiment (treatment:day interaction *F*=15.39, *p*<0.001; Fig. 4B). Despite the latter, co-exposed populations had similar host population dynamics as the populations exposed just to *O. pajunii* (Fig. 2B). All in all, our assumption that *O. pajunii* could be beneficial for *Daphnia* is not supported because *O. pajunii* had a strong negative population-level effect whereas *M. bicuspidata* did not. Citing Francis Bacon “*the remedy is worse than the disease*”.

**Fig. 4.**
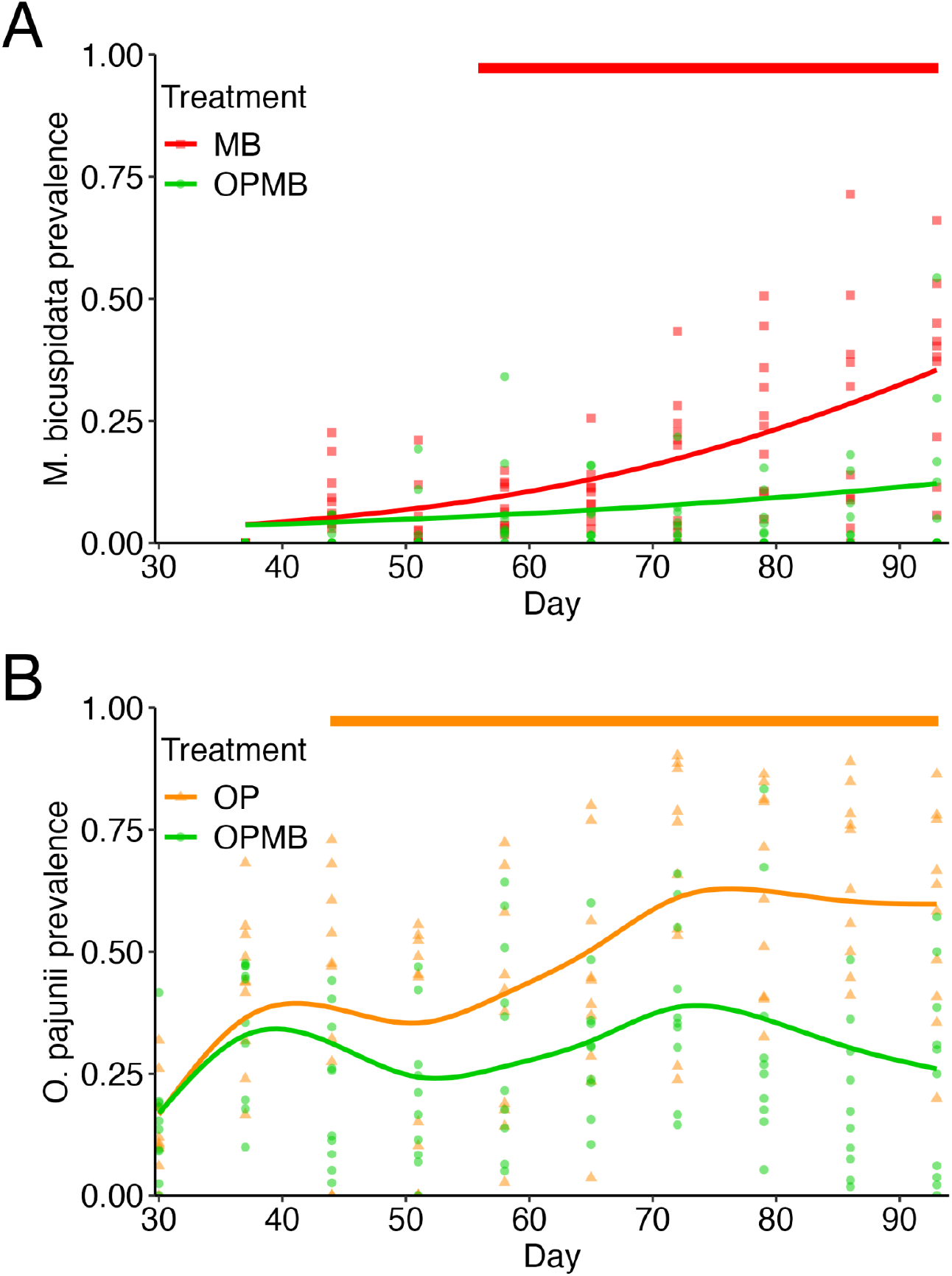
Prevalence of *M. bicuspidata* (A) and *O. pajunii* (B) were lower in the co-infection treatment in comparison to the single infection treatments. The plots show fitted curves of beta regression for *M. bicuspidata* (A) and GAM for *O. pajunii* (B), with data points plotted underneath; the thick lines above the figures show the time span when the prevalence of the respective parasite was higher in the single infection treatment than in the co-infection treatment.

### Beyond individual-level virulence measurements

Awareness of how ecological context affects the outcome of symbiosis is crucial for epidemiology, environmental management and conservation, medicine, and agriculture (*14*). While there is growing consideration of context-dependent changes in symbiosis outcomes, few studies consider the role that time span and level of biological organization play in the functioning of symbiosis. Switching from individual-level analysis to population-level studies done over multiple generations can dramatically change the perspective on the outcome or even nature of the focal symbiosis. For example, anthelmintic treatment that was predicted to help the buffalo population fight off tuberculosis infection actually increased the spread of bovine tuberculosis (*12*). Furthermore, *Wolbachia* is known to have complex transgenerational effects (*40*); for example, in *Drosophila melanogaster*, grandmother age determines the strength of the cytoplasmic incompatibility impact on fitness of the offspring of *Wolbachia*-infected males (*41*). These examples, and many other studies (e.g., (*23, 42–44*)), show that individual-level, single generation studies are burdened with a risk of overlooking important patterns and interactions, such as transgenerational inheritance, selection of host and symbiont genotypes, or shifts in symbiosis outcomes stemming from population/community level interactions. These effects have the potential to change our classification of a focal symbiosis from mutualistic to parasitic or vice versa. Here, we present how transgenerational virulence inflicted by a microsporidian that does minimal harm to the infected host can decimate the host population, and how a highly virulent yeast parasite that was expected to be extremely harmful to the host population turned out to be benign. We would not have been able to predict such strong population-level consequences if we had relied solely on individual-level measurements of virulence in a single generation.

Understanding the large-scale and long-term consequences of symbiosis is crucial for ecology, evolutionary biology, and epidemiology. The strong population-level effects of *O. pajunii* detected in this study indicate that long-lasting epidemics of this widespread microsporidian (as have been seen in nature; (*26, 45, 46*)) have the potential to decrease the abundance of *Daphnia*; given their central role in lake food webs, this likely has consequences for primary producers and ecosystem functioning. Indeed, our finding that there was excess algae in populations exposed to *O. pajunii* but not in control populations or those exposed to only *M. bicuspidata* suggest cascading effects on the food web. More generally, we found that switching between individual- and population-level studies revealed patterns that change our understanding of how a focal symbiosis functions and how it should be classified. Our study emphasizes that individual-based methods of virulence estimation are not enough to make accurate predictions about the impact of a symbiont on a host population, and hence parametrizing models using only within-generation, individual-level measurements may not be sufficient to predict the population- and food web-level effects of epidemics. To combat the spread and harmful effects of infectious diseases in humans, wildlife and agriculture (e.g., (*47–49*)), and to use symbiosis in disease biocontrol (e.g., (*50, 51*)), we require a deep understanding of the outcomes of symbioses at levels beyond individuals. Such understanding comes from determining the impacts of symbionts at the population scale over longer time scales, as well as the role of ecological context in shaping those interactions. Incorporating these factors into our investigations of infectious diseases and symbioses will be crucial not only for furthering our knowledge of the ecology and evolution of symbionts but also for decision making in agriculture, conservation, and management.

## Funding

This work was supported by NSF DEB-1748729 to MAD and by the Gordon and Betty Moore Foundation (GBMF9202) to MAD.

## Author contributions

Conceptualization: MKD, KMM, MHC, MAD

Methodology: MKD, KMM, MHC, MAD

Investigation: MKD, KMM, ESD, FEC, TN, PM, RTM, EB, NFS, MR, ED, KJM, CH, LS, KD

Visualization: MKD

Funding acquisition: MAD

Project administration: MKD, MAD

Supervision: MKD, MAD

Writing – original draft: MKD, MHC, MAD

Writing – review & editing: MKD, ESD, RTM, KJM, MHC, MAD

## Competing Interests

There are no competing interests.

## Data and Materials availability

Data and code used in this manuscript is publicly accessible on GitHub (https://github.com/marcinkdziuba/Pitcher-experiment/releases/tag/v1.1) and through Zenodo (https://doi.org/10.5281/zenodo.15186214).

## Notes

https://doi.org/10.5281/zenodo.15186214

### Competing Interest Statement

The authors have declared no competing interest.

### Summary of Updates

The main text was condensed to fit journal requirements, the algae concentration analysis was changed, an analysis of time at reproduction in individual-level experiment was added to the supplementary information

## References

1. J. Sapp, The dynamics of symbiosis: an historical overview. Can. J. Bot. 82, 1046–1056 (2004).

2. T. L. F. Leung, R. Poulin, Parasitism, commensalism, and mutualism: exploring the many shades of symbioses. Vie Milieu Life Environ. 58,107–115 (2008).

3. A. Casadevall, L. Pirofski, Host-pathogen interactions: the attributes of virulence. J. Infect. Dis. 184, 337–344 (2001).

4. S. R. Thomas, J. S. Elkinton, Pathogenicity and virulence. J. Invertebr. Pathol. 85, 146– 151 (2004).

5. C. E. Cressler, D. V. McLeod, C. Rozins, J. Van Den Hoogen, T. Day, The adaptive evolution of virulence: a review of theoretical predictions and empirical tests. Parasitology 143, 915–930 (2016).

6. R. Poulin, C. Combes, The concept of virulence: Interpretations and implications. Parasitol. Today 15, 474–475 (1999).

7. F. Ben-Ami, J. Routtu, The expression and evolution of virulence in multiple infections: the role of specificity, relative virulence and relative dose. BMC Evol. Biol. 13, 97 (2013).

8. B. A. Hidalgo, L. M. Silva, M. Franz, R. R. Regoes, S. A. O. Armitage, Decomposing virulence to understand bacterial clearance in persistent infections. Nat. Commun. 13, 5023 (2022).

9. Casadevall Arturo, Pirofski Liise-anne, Host-pathogen interactions: Redefining the basic concepts of virulence and pathogenicity. Infect. Immun. 67, 3703–3713 (1999).

10. T. Day, On the evolution of virulence and the relationship between various measures of mortality. Proc. R. Soc. Lond. B Biol. Sci. 269, 1317–1323 (2002).

11. M. Duhamel, P. Vandenkoornhuyse, Sustainable agriculture: possible trajectories from mutualistic symbiosis and plant neodomestication. Trends Plant Sci. 18, 597–600 (2013).

12. V. O. Ezenwa, A. E. Jolles, Opposite effects of anthelmintic treatment on microbial infection at individual versus population scales. Science 347, 175–177 (2015).

13. C. M. Lively, The ecology of virulence. Ecol. Lett. 9, 1089–1095 (2006).

14. J. H. Daskin, R. A. Alford, Context-dependent symbioses and their potential roles in wildlife diseases. Proc. R. Soc. B Biol. Sci. 279, 1457–1465 (2012).

15. T. E. Stewart Merrill, C. E. Cáceres, S. Gray, V. R. Laird, Z. T. Schnitzler, J. C. Buck, Timescale reverses the relationship between host density and infection risk. Proc. R. Soc. B Biol. Sci. 289, 20221106 (2022).

16. R. Poulin, F. Thomas, Epigenetic effects of infection on the phenotype of host offspring: parasites reaching across host generations. Oikos 117, 331–335 (2008).

17. N. C. Johnson, J.-H. Graham, F. A. Smith, Functioning of mycorrhizal associations along the mutualism–parasitism continuum*. New Phytol. 135, 575–585 (1997).

18. C. W. Russell, B. A. Fleming, C. A. Jost, A. Tran, A. T. Stenquist, M. A. Wambaugh, M. P. Bronner, M. A. Mulvey, Context-dependent requirements for FimH and other canonical virulence factors in gut colonization by extraintestinal pathogenic Escherichia coli. Infect. Immun. 86, 10.1128/iai.00746-17 (2018).

19. M. Callens, E. Macke, K. Muylaert, P. Bossier, B. Lievens, M. Waud, E. Decaestecker, Food availability affects the strength of mutualistic host–microbiota interactions in Daphnia magna. ISME J. 10, 911–920 (2016).

20. D. Ebert, Ecology, Epidemiology, and Evolution of Parasitism in Daphnia. (National Library of Medicine (US), 2005).

21. D. Ebert, M. Lipsitch, K. L. Mangin, The effect of parasites on host population density and extinction: experimental epidemiology with Daphnia and six microparasites. Am. Nat. 156, 459–477 (2000).

22. M. A. Duffy, S. R. Hall, Selective predation and rapid evolution can jointly dampen effects of virulent parasites on Daphnia populations. Am. Nat. 171, 499–510 (2008).

23. T. E. Stewart Merrill, S. R. Hall, L. Merrill, C. E. Cáceres, variation in immune defense shapes disease outcomes in laboratory and wild Daphnia. Integr. Comp. Biol. 59, 1203– 1219 (2019).

24. N. R. M. de Albuquerque, K. L. Haag, P. D. Fields, A. Cabalzar, F. Ben-Ami, J.-F. Pombert, D. Ebert, A new microsporidian parasite, Ordospora pajunii sp. nov (Ordosporidae), of Daphnia longispina highlights the value of genomic data for delineating species boundaries. J. Eukaryot. Microbiol. 69, e12902 (2022).

25. M. K. Dziuba, K. M. McIntire, K. Seto, E. S. Davenport, M. A. Rogalski, C. D. Gowler, E. Baird, M. Vaandrager, C. Huerta, R. Jaye, F. E. Corcoran, A. Withrow, S. Ahrendt, A. Salamov, M. Nolan, S. Tejomurthula, K. Barry, I. V. Grigoriev, T. Y. James, M. A. Duffy, Phylogeny, morphology, virulence, ecology, and host range of Ordospora pajunii (Ordosporidae), a microsporidian symbiont of Daphnia spp. mBio 15, e00582–24 (2024).

26. M. A. Rogalski, T. S. Merrill, C. Gowler, C. Caceres, M. Duffy, Context-dependent host-symbiont interactions: shifts along the parasitism–mutualism continuum. Am. Nat. 198, 563–575 (2021).

27. M. K. Dziuba, K. M. McIntire, E. S. Davenport, E. Baird, C. Huerta, R. Jaye, F. Corcoran, P. McCreadie, T. Nelson, M. A. Duffy, Microsporidian coinfection reduces fitness of a fungal pathogen due to rapid host mortality. mBio 15, e00583–24 (2024).

28. K. M. McIntire, M. K. Dziuba, E. B. Haywood, M. L. Robertson, M. Vaandrager, E. Baird, F. Corcoran, M. H. Cortez, M. A. Duffy, Transgenerational pathogen effects: Maternal pathogen exposure reduces offspring fitness. bioRxiv 2023.03.14.532659 [Preprint] (2025). 10.1101/2023.03.14.532659.

29. C.-C. Wei, P.-L. Yen, A. Chaikritsadakarn, C.-W. Huang, C.-H. Chang, V. H.-C. Liao, Parental CuO nanoparticles exposure results in transgenerational toxicity in Caenorhabditis elegans associated with possible epigenetic regulation. Ecotoxicol. Environ. Saf. 203, 111001 (2020).

30. C. U. Braz, T. Taylor, H. Namous, J. Townsend, T. Crenshaw, H. Khatib, Paternal diet induces transgenerational epigenetic inheritance of DNA methylation signatures and phenotypes in sheep model. PNAS Nexus 1, pgac040 (2022).

31. M. D. Klein, A. Proaño, S. Noazin, M. Sciaudone, R. H. Gilman, N. M. Bowman, Risk factors for vertical transmission of Chagas disease: A systematic review and meta-analysis. Int. J. Infect. Dis. 105, 357–373 (2021).

32. J. L. Hite, A. C. Pfenning, C. E. Cressler, Starving the enemy? Feeding behavior shapes host-parasite interactions. Trends Ecol. Evol. 35, 68–80 (2020).

33. S. K. J. R. Auld, K. Raidma, Host genetic variation in feeding rate mediates a fecundity cost of parasite resistance in a Daphnia-parasite system. bioRxiv 2022.11.29.518345 [Preprint] (2022). 10.1101/2022.11.29.518345.

34. E. Davenport, M. Dziuba, F. Corcoran, N. F. dos Santos, K. Monell, P. McCreadie, S. Calhoun, T. Nelson, L. Jacobson, R. Manuel, M. Duffy, Resource quantity affects infection success and impacts of a microsporidian on hosts. Authorea 10.22541/au.173858053.37915414/v1 [Preprint] (2025). 10.22541/au.173858053.37915414/v1.

35. C. L. Searle, M. H. Cortez, K. K. Hunsberger, D. C. Grippi, I. A. Oleksy, C. L. Shaw, S. B. de la Serna, C. L. Lash, K. L. Dhir, M. A. Duffy, Population density, not host competence, drives patterns of disease in an invaded community. Am. Nat. 188, 554–566 (2016).

36. R. M. Penczykowski, M. S. Shocket, J. H. Ochs, B. C. P. Lemanski, H. Sundar, M. A. Duffy, S. R. Hall, Virulent disease epidemics can increase host density by depressing foraging of hosts. Am. Nat. 199, 75–90 (2022).

37. Z. M. Gliwicz, Food thresholds and body size in cladocerans. Nature 343, 638–640 (1990).

38. J. B. McGraw, H. Caswell, Estimation of individual fitness from life-history data. Am. Nat. 147, 47–64 (1996).

39. F. Manzi, S. Halle, L. Seemann, F. Ben-Ami, J. Wolinska, Sequential infection of Daphnia magna by a gut microsporidium followed by a haemolymph yeast decreases transmission of both parasites. Parasitology 148, 1566–1577 (2021).

40. A. R. Weeks, K. Tracy Reynolds, A. A. Hoffmann, Wolbachia dynamics and host effects: what has (and has not) been demonstrated? Trends Ecol. Evol. 17, 257–262 (2002).

41. E. M. Layton, J. On, J. I. Perlmutter, S. R. Bordenstein, J. D. Shropshire, Paternal grandmother age affects the strength of Wolbachia-induced cytoplasmic incompatibility in Drosophila melanogaster. mBio 10, 10.1128/mbio.01879-19 (2019).

42. M. A. Duffy, J. H. Ochs, R. M. Penczykowski, D. J. Civitello, C. A. Klausmeier, S. R. Hall, Ecological context influences epidemic size and parasite-driven evolution. Science 335, 1636 (2012).

43. J. L. Sachs, M. O. Ehinger, E. L. Simms, Origins of cheating and loss of symbiosis in wild Bradyrhizobium. J. Evol. Biol. 23, 1075–1089 (2010).

44. C. C. Wendling, J. Lange, H. Liesegang, M. Sieber, A. Poehlein, B. Bunk, J. Rajkov, H. Goehlich, O. Roth, M. A. Brockhurst, Higher phage virulence accelerates the evolution of host resistance. Proc. R. Soc. B Biol. Sci. 289, 20221070 (2022).

45. E. S. Davenport, M. K. Dziuba, L. E. Jacobson, S. K. Calhoun, K. J. Monell, M. A. Duffy, How does parasite environmental transmission stage concentration change before, during, and after disease outbreaks? Ecology 105, e4235 (2024).

46. E. S. Davenport, M. K. Dziuba, L. E. Jacobson, S. K. Calhoun, K. J. Monell, M. A. Duffy, Parasite transmission stage abundance varies in lakes over time and space. Limnol. Oceanogr. 69, 2167–2179 (2024).

47. E. Tambo, E. C. Ugwu, J. Y. Ngogang, Need of surveillance response systems to combat Ebola outbreaks and other emerging infectious diseases in African countries. Infect. Dis. Poverty 3, 29 (2014).

48. J. Chen, L. Sun, Y. Cheng, Z. Lu, K. Shao, T. Li, C. Hu, H. Han, Graphene oxide-silver nanocomposite: novel agricultural antifungal agent against Fusarium graminearum for crop disease prevention. ACS Appl. Mater. Interfaces 8, 24057–24070 (2016).

49. P. Daszak, J. K. Olival, H. Li, A strategy to prevent future epidemics similar to the 2019-nCoV outbreak. Biosaf. Health 02, 6–8 (2020).

50. L. C. Lewis, D. J. Bruck, J. R. Prasifka, E. S. Raun, Nosema pyrausta: Its biology, history, and potential role in a landscape of transgenic insecticidal crops. Biol. Control 48, 223–231 (2009).

51. D. Sassera, S. Epis, M. Pajoro, C. Bandi, Microbial symbiosis and the control of vector-borne pathogens in tsetse flies, human lice, and triatomine bugs. Pathog. Glob. Health 107, 285–292 (2013).

52. S. N. Wood, Fast stable restricted maximum likelihood and marginal likelihood estimation of semiparametric generalized linear models. J. R. Stat. Soc. Ser. B Stat. Methodol. 73, 3– 36 (2011).

53. R. Lenth, emmeans: Estimated marginal means, aka least-squares means, R package version 1.10.3, GitHub (2024); https://rvlenth.github.io/emmeans/.

54. A. Dinno, _dunn.test: Dunn’s test of multiple comparisons using rank sums_, R package version 1.3.6, CRAN (2024). https://CRAN.R-project.org/package=dunn.test.

55. J. Pinheiro, D. Bates, _nlme: Linear and nonlinear mixed effects models_. R package version 3.1-164, CRAN (2023). https://svn.r-project.org/R-packages/trunk/nlme/

56. F. Cribari-Neto, A. Zeileis, Beta regression in R. J. Stat. Softw. 34, 1–24 (2010).

57. H. Wickham, Ggplot2: elegant graphics for data analysis (Springer-Verlag New York, 2016).

