## Supplementary Materials for "Symbiont virulence is a poor predictor of impacts on host population dynamics"

**The PDF file includes:**

Materials and Methods
Figs. S1 to S2
Table S1

### Materials and Methods

Study system

In this study we exposed *Daphnia dentifera* to two symbionts: *Metschnikowia bicuspidata* and *Ordospora pajunii*. The yeast *M. bicuspidata* is an obligate killer that is ingested by the host during feeding, then penetrates the host's gut to get inside the body cavity, where it proliferates (*23*). Infected host lifespan is reduced (c.a. 12-17 days at 20°C, Fig. 1), and parasite spores are only released after host death (*20*). The spores of the microsporidian *O. pajunii* are also consumed with food, but once they are present in the host's gut lumen, they invade the gut epithelial cells where they reproduce, eventually causing the cell burst and releasing spores back into both the gut lumen and the environment (*25*). As this process is not lethal, *O. pajunii* can reinfect the same host multiple times, which leads to perpetual production and shedding of parasite spores. *O. pajunii* is considered a symbiont of very low virulence, as it has little to no effect on host lifespan or reproduction: Fig. 1 indicates reduction in offspring production only in late life stages – around and past the 5th clutch – and no significant impact on lifespan, while previous studies indicate that it can be completely benign or even beneficial (*25–27*). However, we also observed substantial early mortality of offspring of the *Daphnia* exposed to *O. pajunii*, suggesting that its virulence is mostly transgenerational rather than intragenerational (*28*).

We carried out two experiments, a life-table (or individual-level) experiment and a population experiment. In both experiments, we used the *D. dentifera* clone “S” isolated from a lake in Barry County (Michigan, USA) and kept in the lab for over two decades, the *M. bicuspidata* strain “Standard” isolated from Baker Lake (Barry County, Michigan, USA) in 2003, and the *O. pajunii* isolate “BDWalsh” collected from Walsh Lake (Washtenaw County, Michigan, USA) in 2021.

The S clone was chosen for this study due to its susceptibility to both symbionts and the substantial prior work on it.

Life-table experiment: experimental methods

To estimate the virulence of both symbionts, we designed a life-table experiment with three treatment groups: C - control *Daphnia* not exposed to either one of the symbionts, MB - *Daphnia* exposed to *M. bicuspidata*, and OP - *Daphnia* exposed to *O. pajunii*. We note that these data overlap with some of the data presented in Dziuba et al. (*27*); specifically, the *M. bicuspidata* treatment (MB) individuals in the current experiment are the same as the 'MB high' treatment animals in that experiment (presented in Figure 2 in (*27*)), and the control group (C) in this experiment was also used in supplementary Figure S3 of that article. Prior to the experiment, maternal lines of *Daphnia* were kept singly for three generations in 100 ml of water from North Lake (Washtenaw County, Michigan, USA) filtered with Pall AE glass microfiber filters, fed three times a week with 20,000 cells/ml of green algae *Ankistrodesmus falcatus* (a saturating level of food), and stored in an incubator set at 20°C and 16:8 light:dark conditions.

The experiment started when we collected 0-24h old *Daphnia* of the third clutch of the (uninfected and unexposed) experimental mothers. We pooled the animals in one dish and distributed haphazardly among 90 beakers filled with 100 ml of filtered lake water, having a single *Daphnia* per beaker. Each beaker was randomly assigned to one of the three treatments (C, MB or OP), resulting in 30 replicates per treatment. *Daphnia* were placed in the incubator (same conditions as described above for the stock cultures) and fed 20,000 cells/ml of the green algae daily.

On the third day, the water in all experimental beakers was replaced, with the volume reduced to 50 mL, and *Daphnia* in each group were exposed to their respective treatment: MB received 250 spores/mL of *M. bicuspidata*, OP received a dose of spores equivalent to a single heavily infected donor (which is c.a. 15,000 spores, see (*27*)) and C received a placebo dose composed of ground-up unexposed *Daphnia*, with each animal receiving the equivalent of 1 unexposed animal. To dose *M. bicuspidata* spores for the MB treatment, we harvested *Daphnia* infected with *M. bicuspidata* from our stock cultures, ground them with a motorized pestle, counted the mature spore concentration under a compound scope at x400 magnification using a hemocytometer, diluted the spores with filtered lake water and dosed each beaker with 12,500 spores. To expose the OP treatment, we screened *Daphnia* from lab *O. pajunii* stock cultures under a dissecting microscope and harvested heavily infected individuals, placed them in a 1.5 ml centrifuge tube filled with 100 µl of filtered lake water, then ground them with a pestle into a slurry. The slurry was resuspended in filtered lake water and dosed into the beakers in a ratio of 1 donor:1 recipient, so each beaker received a volume of slurry equivalent to a single ground-up donor *Daphnia*. We did not count the *O. pajunii* spores due to their small size (~2µm length), which makes it impossible to determine whether a given spore is mature/infective or not. To ensure that each beaker received a dose of similar composition, we dosed all the beakers from the same spore slurry, as we have done in other experiments (e.g., (*27*)). We prepared the placebo solution by collecting uninfected *Daphnia* from cultures that had never been exposed to *O. pajunii* in a 1.5 ml centrifuge tube, grinding them with a motorized pestle and resuspending the slurry in filtered lake water; we dosed the C treatment beakers with the placebo slurry in a ratio of one ground-up *Daphnia* per beaker. The exposure lasted two days for all treatments, after which we changed water and beakers for each *Daphnia*, and carried out the experiment until all *Daphnia* had died. We monitored the *Daphnia* daily for survival and offspring production, and we counted and removed the offspring each day. We checked each parasite-exposed *Daphnia* for infection at death or on day 20, whichever came first (20 days is enough for individuals exposed to either pathogen to show visible symptoms of infection). For both symbionts, 100% of the exposed animals who lived long enough for infections to develop were infected.

<u>Life table experiment: statistical analysis</u>

To compare the virulence of both symbionts, we analyzed the number of offspring produced by *Daphnia* per clutch (reproductive event) and the host's lifespan in each treatment. For the reproduction analysis, we used a generalized additive model (GAM) from the package *mgcv* (*52*) because, as *Daphnia* grow, their clutch size increases, but this increase eventually stops and *Daphnia* start producing fewer offspring due to senescence; hence the change of clutch size over time is non-linear. We also trimmed out the data past clutch fourteen because the estimates would have been unreliable due to high mortality and low number of replicates later in the experiment. We used number of offspring as the response variable, parasite treatment, clutch number, and their interaction as explanatory variables, and clutch number as a common smoother for all the treatments. Based on a comparison of Akaike Information Criterion (AIC), we determined that there was no justification to use separate smoothers for each treatment. The model was investigated for normality of residuals distribution, homogeneity of variances, and influential outliers using diagnostic plots. Post-hoc pairwise comparisons were conducted with estimated marginal means and Bonferroni correction using the R package *emmeans* (*53*). The GAM model diagnostics showed a slight increase in the response variable plotted against the fitted values because the first few *Daphnia* clutches naturally have fewer offspring than later clutches (due to small size of the mothers). To be sure that this small inconsistency in variance does not make our results qualitatively erroneous, we conducted an additional set of non-parametric Kruskal-Wallis tests of the relationship between the number of offspring per clutch and parasite treatment at each clutch, followed by a Dunn test from package *dunn.test* (*54*) for pairwise comparisons. The results of this test were consistent with the GAM results, although a bit more conservative; we do not show the non-parametric test results in the manuscript, but they are available in the R code, which is publicly accessible on GitHub (https://github.com/marcinkdziuba/Pitcher-experiment/releases/tag/v1.1) and through Zenodo (https://doi.org/10.5281/zenodo.15186214).

Mean lifespan of *Daphnia* was compared between the treatments using a generalized least squares model with lifespan (measured in days) as the response variable, parasite treatment as the explanatory variable and as a variance covariate due to heterogeneity of variances between the treatments. We checked model assumptions with diagnostic plots and conducted pairwise comparisons with *emmeans* and using Bonferroni correction.

Age at reproduction of *Daphnia* during the first four clutches was compared between treatments using linear mixed effects model from *nlme* package (*55*) with age (in days) as the response variable, parasite treatment as the explanatory variable and as a variance covariate due to heterogeneity of variances between the treatments, and clutch as a random intercept. After checking model assumptions with diagnostic plots, we conducted pairwise comparisons with *emmeans* and using Bonferroni correction.

<u>Population experiment: experimental methods</u>

We designed this experiment to test the effects of *M. bicuspidata* and *O. pajunii* on *Daphnia* populations, as well as the protective effect of *O. pajunii* against *M. bicuspidata* outbreaks. We used the same genotypes of host and symbionts for this experiment as for the life-table experiment. The maternal lines of *Daphnia* were prepared following the same protocol as the life table experiment. Our population-level experiment started when the (uninfected and unexposed) mothers produced their second clutch of offspring, which we collected 0-24h after birth. We pooled the offspring in one dish and distributed them haphazardly among 40 beakers filled with 100 ml of filtered lake water, adding one individual per beaker, then a second to each beaker and so on until each beaker had 10 juvenile *Daphnia*. We assigned each beaker to one of four treatments (10 replicates each): C - control treatment with only *Daphnia* and no parasites; MB - *Daphnia* exposed to *M. bicuspidata* spores on day 31; OP - *Daphnia* exposed to *O. pajunii* spores on day 1; OPMB - *Daphnia* exposed to *O. pajunii* spores on day 1 and then to *M. bicuspidata* spores on day 31. After assigning the beakers to treatments, we exposed OP and OPMB treatments to spores of *O. pajunii*. The spore slurries were dosed the same way as in the life table experiment, except this time the dosing ratio was 1 donor : 2 recipients, so each beaker containing 10 juvenile *Daphnia* received a volume of slurry equivalent to 5 ground-up donor *Daphnia*. After dosing, all treatments were fed 20,000 cells/ml of algae and placed in an experimental room maintained at 20°C and with 16:8 light:dark conditions. After two days, we poured the content of each beaker into a separate 5L pitcher filled with 5L of filtered lake water, in which *Daphnia* populations were kept for the rest of the experiment. Each 5L pitcher was dosed with 10,000 cells/ml of algae on this initial transfer day and then three times a week throughout the experiment.

For each pitcher, we took weekly measurements of *Daphnia* density, infection prevalence, demographic structure (females, males, juveniles), and algae concentration, beginning on day 9 of the experiment. The screening protocol was as follows: the content of each pitcher was transferred into a clean 5L pitcher (to prevent periphyton buildup), the water was refilled when needed to account for evaporative losses, the contents of the pitcher were gently stirred using a 150 ml ladle to homogenize *Daphnia* distribution across the water column, and then subsampled with the ladle. The subsamples were at least 300 ml (with an exception of day 30 when we subsampled 150 ml as *Daphnia* densities were extremely high) and had at least 20 *Daphnia*. We did not subsample more than 1000 ml. In the case of densities below detection, we quantified the number of *Daphnia* in the whole pitcher. A subsample was poured into a 2L watch glass, and each *Daphnia* from the subsample was then transferred with a glass pipette onto a microscope slide and checked under a dissecting microscope for infection and demographic identity. Processed *Daphnia* were returned to their pitchers. After checking all *Daphnia*, we measured the concentration of green algae in 50 ml of water using a FluoroProbe (BBE Moldanke). The sample to determine algal standing stock was always collected on the third day after the previous feeding and right before the next feeding. The rest of the water from the subsample was poured from the watch glass back into the respective pitcher. Each treatment had a separate ladle, and the glassware was changed between each treatment to avoid spore transfer between treatments.

Screening on day 30 revealed that *O. pajunii* epidemics were established, with ~15% parasite prevalence in the exposed populations. Therefore, on day 31, treatments MB and OPMB were exposed to *M. bicuspidata* spores. The *M. bicuspidata* dosing was staggered because we were testing the hypothesis that *M. bicuspidata* is less successful when the host is already infected with

*O. pajunii* (*27*). Additionally, our field data indicated that *O. pajunii* epidemics occur before *M. bicuspidata* outbreaks (*27*). Thus, we needed the *O. pajunii* epidemics to be established before introducing *M. bicuspidata*. We prepared the *M. bicuspidata* slurries following the same protocol as the life table experiment. However, we dosed the pitchers with 12.5 spores/ml of water (62,500 spores per pitcher), which is lower than the dose used in the individual-level experiment (i.e., 250 spores/ml), because previous studies showed that higher experimental volumes required lower spore concentrations to yield high infection levels (*35*). After dosing the pitchers with *M. bicuspidata*, we continued screening the 40 populations for prevalence of both parasites and the other parameters until day 93 of the experiment. We terminated the experiment at day 93 because some of the beakers started growing filamentous algae in which the *Daphnia* were getting tangled and stuck, leading to their starvation and death. We note that, in the OPMB treatment, very advanced and severe *M. bicuspidata* infections could have obscured the diagnosis of *O. pajunii* infections later in the experiment. Spores of *M. bicuspidata* in heavily infected hosts overgrow the hemocoel around the gut, blocking our view of the gut. Therefore, there is a risk that some animals that were actually co-infected in the OPMB treatment late in the experiment were scored as singly infected with only *M. bicuspidata*, which could have resulted in underestimation of *O. pajunii* prevalence in the OPMB treatment.

Population experiment: statistical analysis

The density of *Daphnia* was analyzed as the total number of all *Daphnia* (infected and uninfected, adults and juveniles, females and males) per liter of water. Based on the density data at the beginning of the experiment and on day 23 – the last screening day of the initial growth phase – we calculated population growth rates using the following equation: $r$=ln(density$_{t23}$ − density$_{t0}$)/n, where $\text{density}_{t23}$ is *Daphnia* density at day 23, $\text{density}_{t0}$ is *Daphnia* density at the beginning of the experiment and n is number of days. Because we analyzed the time period prior to *M. bicuspidata* spore dosing, we effectively had only two groups of populations: exposed to *O. pajunii* OP (containing treatments OP and OPMB) and unexposed C (containing treatments C and MB), and we analyzed the data as such. Due to influential outliers and violation of normal distribution of residuals, we analyzed the data with a non-parametric Kruskall-Wallis test.

To investigate the change in population size over time (after day 30), we used a GAM with *Daphnia* density as a response variable, treatment, day of experiment, and their interaction as explanatory variables, and day as a smoothing factor fitted separately for each treatment. After fitting the model, we inspected diagnostic plots to ensure the assumptions of normal distribution of residuals and homogeneity of variance were met. Due to significant interactions between the terms, we conducted a pairwise comparison with Bonferroni correction to determine at which days population densities are significantly different, using the package *emmeans*.

To analyze *M. bicuspidata* prevalence in treatments MB and OPMB, we used a beta regression model from the package *betareg* (*56*) with the day of experiment, treatment, and their interaction as explanatory variables. We used beta regression because *M. bicuspidata* prevalence data were strongly right-skewed (many measurements close to zero), and beta regression handles the skewed distribution of proportion data much better than a GAM. To analyze the prevalence of *O. pajunii* in groups OP and OPMB, we used a GAM with day of experiment, treatment, and their interaction as explanatory variables, and day as a smoothing factor common for both groups (based on AIC score for common vs. separate smoothers). For both prevalence analyses, we conducted model diagnostics and used the *emmeans* package and Bonferroni correction to determine at which days the prevalence differed between the compared treatments.

The proportion of juveniles (juvenile density divided by total density) was analyzed using a linear model with day of experiment, treatment, and their interaction as explanatory variables, checking model diagnostics and conducting pairwise comparisons using *emmeans* and Bonferroni correction.

The data on algae concentration in the pitchers had a large proportion of zeros (indicating that *Daphnia* in those populations consumed all the algae provided). Thus, we used a GAM with treatment as the explanatory variable, day as a smoothing factor fitted separately for each treatment, and a Tweedie distribution with log link. After fitting the model, we confirmed homogeneity of variances and conducted pairwise comparisons with *emmeans* and Bonferroni correction to determine at which days the algal density differed between the parasite-exposed treatments and the control. All analyses and figures were conducted in RStudio using R version 4.4.1, and the figures were plotted using the package *ggplot2* (*57*).

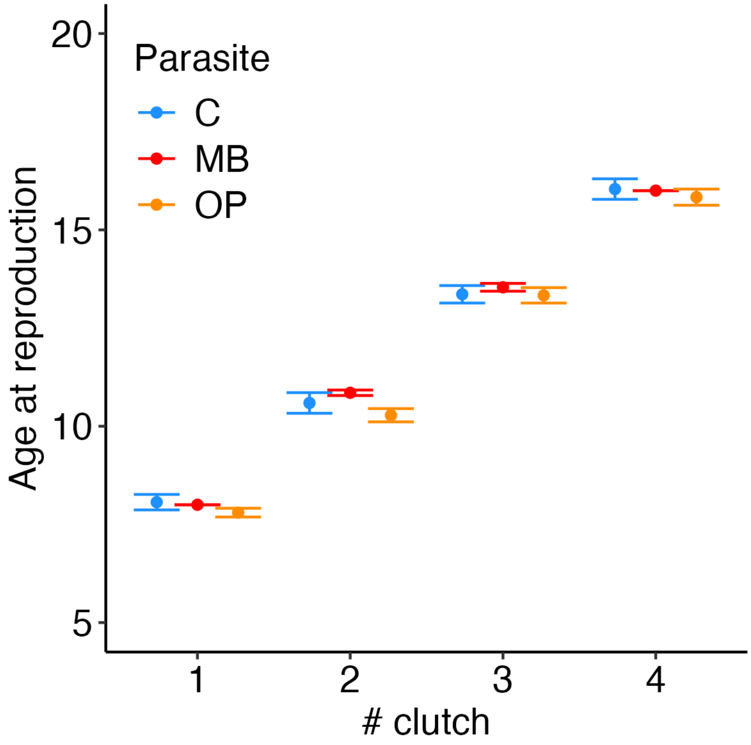


**Fig. S1.**

***Daphnia* exposed to *M. bicuspidata* reproduced later than controls and *O. pajunii* exposed *Daphnia* at clutches 2 and 3, while *Daphnia* exposed to *O. pajunii* reproduced at the same time as controls in first four clutches.** Mean±SE age at reproduction in clutches 1-4 is presented for *Daphnia* exposed to *M. bicuspidata*, *O. pajunii* and unexposed controls.

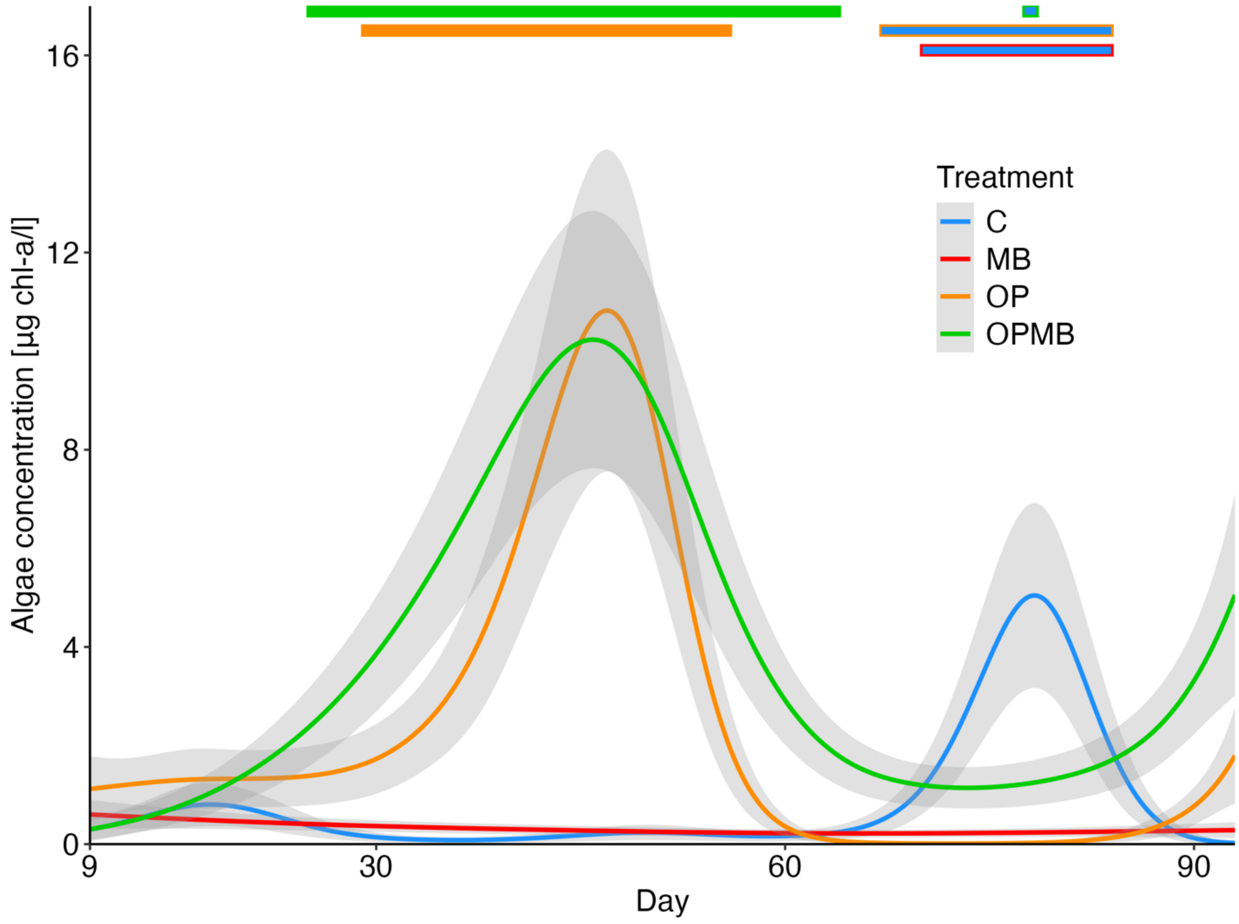


**Fig. S2.**

**Algae were mostly depleted in control and *M. bicuspidata*-exposed treatments, but treatments with *O. pajunii* outbreaks had leftover algae present in time overlapping with lowered *Daphnia* density (Fig. 2).** The OP and OPMB treatments had excess algae on days 29-56 and 25-64, respectively. Control treatment C had significantly more algae than MB on days 70-84. The solid lines and grey ribbons indicate GAM modeled fitted values±SE of algae concentration in pitchers over time. Thick orange and green lines on top of the figure indicate the time span at which OP and OPMB, respectively, had significantly larger algae concentration than control; the blue lines with red, orange and green borders indicate the time span at which control treatment C had higher algae concentration in comparison to MB, OP and OPMB, respectively.

**Table S1.**

**Comparison of *Daphnia* age at reproduction at first four clutches of the individual-level experiment.** The comparison was conducted with Kruskal-Wallis test, and when the comparison yielded significant difference, it was followed by a pairwise Dunn test.

| Clutch | K-W χ2 | K-W p-value | pair | Dunn p-value |
|---|---|---|---|---|
| 1 | 4.9355 | 0.085 | | |
| 2 | 20.633 | <0.001 | C-MB | <0.001 |
| 2 | | | C-OP | 0.623 |
| 2 | | | MB-OP | <0.001 |
| 3 | 9.0596 | 0.011 | C-MB | 0.012 |
| 3 | | | C-OP | 1 |
| 3 | | | MB-OP | 0.018 |
| 4 | 1.0361 | 0.596 | | |
